# Magnetic Field–Enhanced Docosahexaenoic Acid Delivery for Acetylcholine Restoration in Alzheimer’s Disease: an In Silico Study

**DOI:** 10.64898/2026.09.28.755051

**Authors:** Madhuleena Dasgupta, Amit Konar, Atulya Nagar

**Affiliations:** School of Mathematics, Computer Science and Engineering, Liverpool Hope University, UK; Artificial Intelligence Lab, ETCE Department, Jadavpur University, Kolkata, India

**Keywords:** Alzheimer’s Disease, Acetylcholine, Docosahexaenoic Acid, Static magnetic field, In silico modelling, Cholinergic restoration

## Abstract

Alzheimer’s Disease (AD) is characterised by the progressive depletion of Acetylcholine (ACh), a neurotransmitter essential for cognition and synaptic communication. Docosahexaenoic acid (DHA), the predominant omega-3 polyunsaturated fatty acid in neuronal membranes, raises cerebral ACh and supports cholinergic transmission, but its therapeutic efficacy is limited by poor blood–brain barrier permeability and slow passive diffusion. This work addresses these limitations through a closed form mathematical model coupling the spatiotemporal dynamics of DHA diffusion and ACh synthesis under an externally applied static magnetic field. Because DHA is diamagnetically anisotropic, it aligns along the field axis through diamagnetic torque while body cells form field-aligned linear arrays; together these effects raise the effective diffusion coefficient of DHA. At a representative observation point, 0.5 cm, magnetic field–assisted transport reduces the time for ACh to reach a therapeutic threshold by 83.3%, from 10.55 s to 1.76 s. This reduction is spatially uniform across the modelled range of 0–3.0 cm from the source, with the absolute saving growing from 6.71 s at the source to 19.21 s at 3.0 cm. Co-evolution analysis reveals the mechanism: by the end of a 2.0 s window the field elevates DHA 7.39-fold relative to baseline, driving ACh to 133.7% of threshold versus only 18.1% without the field. A Brain Computer Interface (BCI)-based closed-loop dosing framework is additionally proposed for future clinical validation. These findings establish a quantitatively grounded theoretical basis for magnetically enhanced DHA therapy as an approach to cholinergic restoration in AD.

## 1 Introduction

Alzheimer’s Disease (AD) is a progressive neurodegenerative disorder characterised by cognitive decline, memory loss, and impaired synaptic communication, affecting millions worldwide. A central hallmark of AD is the depletion of acetylcholine (ACh), a neurotransmitter essential for cognitive processing and synaptic function (Kasa *et al*. 1997; Francis *et al*. 1999). Current cholinesterase inhibitors such as donepezil, rivastigmine, and galantamine provide temporary symptomatic relief. However, they do not restore physiological ACh levels or halt neurodegenerative progression (Birks and Harvey 2018; Yiannopoulou and Papageorgiou 2020; Cummings *et al*. 2021). This motivates the search for strategies that address the underlying cholinergic deficit more directly.

The brain is enriched with long-chain omega-3 polyunsaturated fatty acids (PUFAs) (Rapoport *et al*. 2011; Weiser *et al*. 2016; Duro *et al*. 2023). Docosahexaenoic acid (DHA) is the predominant omega-3 PUFA. It constitutes a major structural component of neuronal membrane phospholipids and is essential for maintaining normal brain structure, function, and metabolism (Rapoport *et al*. 2011; Weiser *et al*. 2016). Neurotransmission depends strongly on membrane phospholipid composition. Neurons enriched in DHA exhibit enhanced vesicle formation and fusion efficiency, influencing neurotransmission fidelity (Vancassel *et al*. 2007). DHA deficiency has accordingly been linked to impaired neurotransmitter release and synaptic transmission.

The dietary precursor of omega-3 PUFA is alpha-linolenic acid (ALA) (Singh 2005). Deficiency of ALA during brain development induces significant changes in neurotransmitter release in cerebral areas involved in learning, memory, and reward (Vancassel *et al*. 2007). Affected neurotransmitters include ACh, dopamine, and serotonin. Additionally, alterations in presynaptic storage processes and disruptions of monoaminergic–cholinergic pathway interactions are observed. With respect to ACh specifically, DHA deficiency produces a 10% decrease in muscarinic receptor binding, although acetylcholinesterase activity and the vesicular ACh transporter are not affected (Aïd *et al*. 2003; Tanaka *et al*. 2012). Conversely, dietary DHA supplementation increases cerebral ACh levels (Minami *et al*. 1997). These findings collectively motivate the use of DHA as a regulator of cholinergic signalling and cognitive recovery in AD.

Despite these promising findings, the therapeutic efficacy of conventional DHA supplementation remains limited. Poor bioavailability, restricted blood–brain barrier (BBB) permeability, and inadequate diffusion across brain tissue collectively reduce DHA availability at target neural sites (Pan *et al*. 2015; Iwao *et al*. 2023; Duro *et al*. 2023; Ali *et al*. 2024). Recent advances in magnetic field–assisted drug delivery have demonstrated promising capabilities for targeting the central nervous system. Senanayake *et al*. (2025) reviewed magnetic targeting strategies that exploit magnetic nanoparticles (MNPs) to overcome BBB limitations, achieving enhanced spatial penetration and controlled drug distribution in neural tissue. Alipour *et al*. (2025) highlighted magnetothermal stimulation, wherein magnetic fields induce localised heating of functionalised nanoparticles, enabling modulation of ion channels and neuronal excitability. These MNP-based approaches, however, require exogenous nanoparticle administration and involve complex preparation protocols. The present work takes a fundamentally different approach: rather than employing nanoparticles, it exploits the intrinsic diamagnetic anisotropy of DHA molecules themselves under an externally applied static magnetic field, representing a simpler, non-invasive strategy for magnetically enhanced DHA delivery.

Given DHA’s high degree of unsaturation and consequent anisotropic molecular structure, it exhibits diamagnetic anisotropy. Under an externally applied static magnetic field, two biophysical mechanisms operate simultaneously. First, DHA molecules align along the field axis through diamagnetic torque, enhancing their directional transport toward the target site. Second, body cells form linear arrays aligned with the field direction, creating a structured pathway that further facilitates DHA diffusion through tissue. Together, these effects enhance the effective diffusion coefficient of DHA, enabling more efficient distribution within target neural regions. This approach is non-invasive, requires no exogenous agents, and draws on the established biophysical principle that diamagnetically anisotropic molecules reorient under a static magnetic field (Rosen 2003).

Despite the biological rationale for DHA-mediated cholinergic enhancement and the physical plausibility of magnetic field–assisted delivery, no closed-form analytical framework currently exists that couples DHA transport physics with ACh synthesis kinetics under static magnetic field modulation. Existing models are either purely pharmacokinetic, addressing DHA metabolism without magnetic enhancement, or focused on nanoparticle-based delivery without cholinergic coupling. The quantitative relationship between magnetic field–enhanced DHA diffusion and the consequent rate of ACh restoration remains uncharacterised. The present work addresses this gap.

### 1.1 Key Neurobiological Terms

For conceptual clarity and consistency, key biological and signal-processing terms used throughout this study are defined below. The first four terms relate to the core biochemical framework. The final two terms relate to the Brain–Computer Interface dosing framework presented in Section 4.4.

#### Acetylcholine (ACh)

A neurotransmitter synthesised in cholinergic neurons that plays a critical role in cognitive processing, memory consolidation, attention, and motor co-ordination.

#### Docosahexaenoic Acid (DHA)

A 22-carbon omega-3 polyunsaturated fatty acid (22:6 n-3) and the predominant structural lipid in neuronal membranes. DHA contributes to membrane fluidity, vesicle fusion efficiency, and neurotransmitter release dynamics, thereby supporting cholinergic signalling integrity.

#### Cholinergic Neurotransmission

The sequence of events by which ACh is synthesised, stored in synaptic vesicles, released upon neuronal activation, and subsequently binds to postsynaptic receptors or undergoes enzymatic degradation by acetylcholinesterase.

#### Exocytosis

The calcium-triggered fusion of neurotransmitter-containing vesicles with the presynaptic membrane, resulting in transmitter release into the synaptic cleft. Membrane lipid composition, including DHA content, strongly influences vesicle fusion probability and release kinetics.

#### Event-Related Potential P300

A positive deflection in the electroencephalographic (EEG) signal occurring approximately 300 ms after presentation of a task-relevant target stimulus. It reflects cognitive evaluation, attention allocation, and stimulus categorisation processes.

#### Event-Related Desynchronization/Synchronization (ERD/ERS)

Transient decreases (ERD) or increases (ERS) in oscillatory cortical rhythms, associated with motor intention, task engagement, and cortical activation patterns.

### 1.2 Problem Statement

Despite extensive evidence supporting the neuroprotective role of DHA in sustaining cholinergic transmission (Kim *et al*. 2022), its therapeutic efficacy in AD remains limited by three interconnected clinical challenges. First, conventional DHA supplementation fails to achieve uniform distribution within neural tissue due to restricted BBB permeability, slow diffusion kinetics, and rapid peripheral metabolism (Pan *et al*. 2015; Hachem *et al*. 2020; Iwao *et al*. 2023; Ali *et al*. 2024). Second, no quantitative model presently exists that predicts how magnetic field modulation accelerates DHA delivery and restores ACh to therapeutic levels as a function of distance from the source. Third, the absence of adaptive feedback mechanisms prevents real-time adjustment of DHA dosing in response to changing neurophysiological states.

To address these challenges, the present study formulates a comprehensive *in silico* framework with the following original contributions:

*Firstly*, a closed-form mathematical framework is developed that couples DHA diffusion and ACh synthesis under an externally applied static magnetic field, yielding closed-form analytical expressions for DHA concentration, ACh concentration, and the time required to reach a therapeutic ACh threshold under both baseline and magnetically enhanced conditions. The DHA expression solves Fick’s second law directly; the ACh concentration and threshold-time expressions are exact for the effective linearised form of the coupled model.

*Secondly*, magnetic field–enhanced DHA diffusion is quantified across the modelled spatial range (0–3.0 cm measured from the DHA source). At a representative observation point (*x = 0*.*5 cm*), the field reduces the ACh threshold attainment time by 83.3% (from 10.55 s to 1.76 s). This proportional reduction is spatially uniform throughout the modelled range, with the absolute time saving growing from 6.71 s at the source to 19.21 s at the most distal point evaluated (3.0 cm). *Thirdly*, co-evolution analysis of DHA and ACh concentrations identifies the underlying mechanism. By the end of a 2.0 s observation window (*t = 2*.*0 s*), magnetic field assistance has elevated DHA availability 7.39-fold relative to baseline at the target site, driving ACh to 133.7% of the therapeutic threshold, whereas baseline conditions reach only 18.1%. This establishes the mechanistic basis of the observed improvement. *Lastly*, a Brain–Computer Interface (BCI)-based closed-loop DHA dose regulation framework is proposed, utilising P300 and ERD/ERS biomarkers to adaptively modulate DHA administration across three physiologically grounded dosing states. The framework is demonstrated through simulation as a pathway for future clinical validation.

The remainder of this paper is organised as follows. Section 2 presents the mathematical framework, including the governing equations for DHA diffusion and ACh synthesis under static magnetic field modulation, together with the model parameter table. Section 3 describes the simulation methodology and algorithmic implementation. Section 4 presents simulation results across three quantitative outputs: ACh concentration dynamics (Section 4.1), spatial threshold analysis (Section 4.2), and DHA-ACh co-evolution (Section 4.3), followed by the BCI-based closed-loop dosing framework (Section 4.4). Section 5 discusses the findings in the context of cholinergic restoration in AD. Section 6 addresses limitations and future experimental directions. Section 7 concludes the paper.

## 2 Mathematical Framework

The mathematical framework models two coupled biological processes. DHA diffuses through brain tissue from the administration site to the target neural region. As it arrives, it drives the synthesis of ACh. The framework yields closed-form analytical expressions for three quantities: the DHA concentration, the ACh concentration, and the time for ACh to reach a therapeutic threshold. These are obtained under both baseline and magnetically enhanced conditions. The derivation proceeds in four stages (Sections 2.1–2.4), followed by the model parameters in Section 2.5.

All quantities in this framework are expressed in normalised, dimensionless form. Spatial position is scaled to a characteristic diffusion length (*λ*_*0*_ *= 1 cm*). Time is scaled to a characteristic timescale (*τ*_*0*_ *= 1 s*). The magnetic field is scaled to a reference field strength (*B*_*ref*_). Under this non-dimensionalisation, the diffusion coefficient *η*, the effective diffusion coefficient *D*_*eff*_, the decay rate *β*, the magnetic proportionality constant *k*_*B*_, and the field magnitude *B* are all dimensionless. Every exponential argument is therefore dimensionless by construction. Distances and times reported later are multiples of *λ*_*0*_ and *τ*_*0*_. For example, *x = 0*.*5* means 0.5 cm, and *t = 1*.*76* means 1.76 s.

### 2.1 Governing Equation of DHA-ACh Coupled Dynamics

The starting point is a simple observation: ACh synthesis in the brain is promoted by the availability of DHA at the postsynaptic membrane. DHA enrichment enhances vesicle formation and fusion efficiency, which augments cholinergic signalling (Minami *et al*. 1997; Vancassel *et al*. 2007). At the same time, ACh is broken down by acetyl-cholinesterase at a rate proportional to its current concentration. Let *x*_0_ denote the source position of DHA and *t*_0_ the initial time. Let *[ACh]*_*(x, t)*_ and *[DHA]*_*(x, t*)_ denote the concentrations of ACh and DHA at distance *x* from the source at time *t*. These two competing processes: DHA-driven synthesis and natural decay are captured by the following first-order ordinary differential equation:

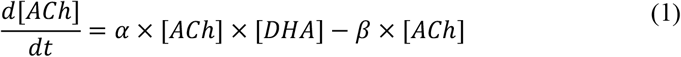

Where *α* is the ACh–DHA biochemical coupling constant. It governs the rate at which DHA availability promotes ACh synthesis. The constant *β* is the natural ACh decay rate. The product term (*α ×[ACh]×[DHA]*) represents ACh synthesis; its rate depends on both the current ACh level and the available DHA. The term (−*β×[ACh]*) represents natural decay through enzymatic degradation.

### 2.2 DHA Diffusion and Closed-Form Concentration Expressions

The distribution of DHA in space and time is governed by Fick’s second law of diffusion:

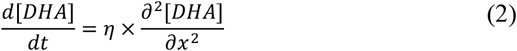

Where *η* is the (normalised) diffusion coefficient. A separable solution of Equation (2) is given below. It combines exponential attenuation in space with accumulation in time:

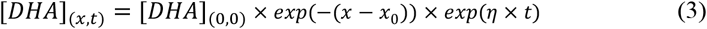

Where *[DHA]*_*(0,0)*_ is the initial DHA concentration at the source. The factor *exp*(−(*x*− *x*_0_)) describes how DHA attenuates with distance from the source. The factor *exp*(η × *t*)describes how DHA accumulates at the observation point over time. Equation (3) can be verified by substitution: both 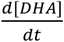 and 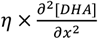equal *η×[DHA]*, so Equation (2) is satisfied.

The temporal factor, *exp*(η × *t*) represents a source-replenished regime, in which DHA is continuously supplied at the administration site. It is intended to hold over the short observation window (0–2 s) analysed here, not as an unbounded steady state. Over this window, it reproduces the early, near-monotonic rise in target-site DHA that drives cholinergic synthesis.

To obtain a tractable closed form for ACh, the DHA-dependent synthesis term in Equation (1) is represented by an effective first-order growth process. Specifically, the synthesis rate *α×[DHA]* is modelled as an effective rate constant equal to the diffusion coefficient, *η*. The coupling constant *α* is thereby absorbed into this effective rate. Under this effective-rate (linearising) closure, Equation (1) reduces at fixed *x* to the linear equation: *d[ACh]/dt* = (η − β) × [*ACh*]. Its exact solution is:

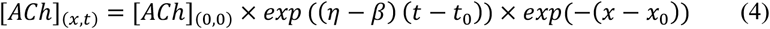

Where, [*ACh*]_(0,0)_ is the initial ACh concentration, and the spatial factor *exp*(−(*x*− *x*_0_)) is inherited from the DHA profile. Equation (4) is therefore an exact closed-form solution of the effective linearised model, not of the full nonlinear system in Equation (1). It is this reduction that makes the subsequent threshold-time analysis analytically exact. The net growth rate (*η*−*β*) governs how ACh evolves over time. When *η* > *β*, ACh increases steadily toward the therapeutic threshold. For the present parameters (*η = 0*.*5, β = 0*.*3*), (*η*−*β*) *= 0*.*2 > 0*, which ensures monotonic ACh growth under baseline conditions.

### 2.3 Time to Therapeutic Threshold

The therapeutic threshold is reached when ACh first equals *[ACh]*_*th*_ *= 5 × [ACh]*_*(0,0)*_. This time is found by setting *[ACh]*_*(x, t)*_ *= [ACh]*_*th*_ in Equation (4) and solving for *t*. The result is the closed-form threshold-time expression:

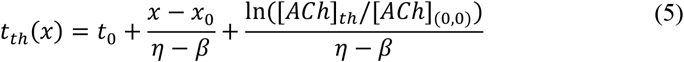

Equation (5) is the principal analytical result of the baseline framework. It gives the exact threshold-attainment time for any observation distance *x*, with no numerical approximation. The distance-dependent term, 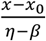, is the travel time of the leading edge of the spreading DHA from source to observation point. It grows linearly with distance, which explains the linear relationship in Fig. 2. The remaining term, 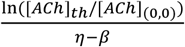 is the time for ACh to grow from its initial level to the threshold. This term does not depend on distance. It depends only on the threshold ratio and the net growth rate.

### 2.4 Magnetic Field-Enhanced DHA Transport

The static magnetic field, *B* is generated by wearable current-carrying coils: a wrist band, a pendant, and a headband following 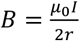,where, *I* is the current through a coil, *r* is its radius, and *μ*_*0*_ is the permeability of the coil material. Here *B* is expressed in normalised form, relative to the reference field *B*_*ref*_. Under this field, two biophysical mechanisms enhance DHA transport. First, DHA molecules are diamagnetically anisotropic, so they align along the field axis through diamagnetic torque (Rosen 2003). This alignment enhances directional transport toward the target site. Second, active body cells form linear arrays aligned with the field. These arrays create a structured pathway that facilitates DHA diffusion through tissue. The combined effect of these two mechanisms is modelled as an enhanced effective diffusion coefficient:

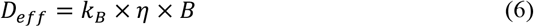

Where, *k*_*B*_ is a dimensionless magnetic proportionality constant. It characterises the combined efficiency of both mechanisms. In the normalised formulation, *η, B*, and *k*_*B*_ are all dimensionless. Their product is therefore dimensionless, so *D*_*eff*_ is dimensionless like *η*, and Equation (6) is dimensionally consistent.

Replacing *η* with *D*_*eff*_ in Equations (3), (4), and (5) gives the corresponding with-field expressions for DHA concentration, ACh concentration, and threshold time:

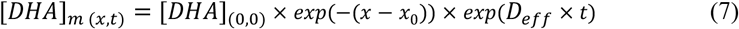

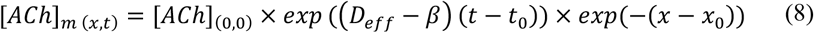

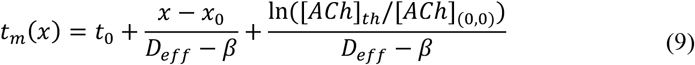

Equations (5) and (9) together form the complete analytical basis for all simulation results in Section 4. The simulation is conducted at an observation point *x = 0*.*5* from the DHA source, with *t*_*0*_ *= 0* and *x*_*0*_ *= 0*.

A useful analytical consequence follows directly. With *t*_*0*_ *= x*_*0*_ *= 0*, both threshold times share the common factor, (*x*+ ln([*ACh*]_*th*_/[*ACh*]_(0,0)_). Their ratio is therefore independent of distance:

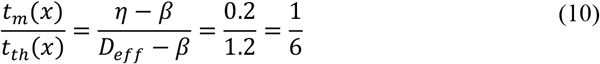

Hence, the magnetic field reduces the threshold-attainment time by exactly 5/6 (83.3%), uniformly across all distances. This is an exact analytical property of the model, not a numerical coincidence. It underlies the spatially uniform reduction reported in Section 4.

### 2.5 Model Parameters

Table 1 lists all model parameters. All values are theoretically motivated. They are chosen to ensure three things: (i) biological plausibility of the concentration profiles, (ii) internal consistency of the closed-form solutions, and (iii) qualitative agreement with published evidence on DHA–ACh interactions (Minami *et al*. 1997; Vancassel *et al*. 2007). The cited references justify the modelling framework and the physical regime. They are not direct sources for the specific numerical values adopted.

**Table 1.**
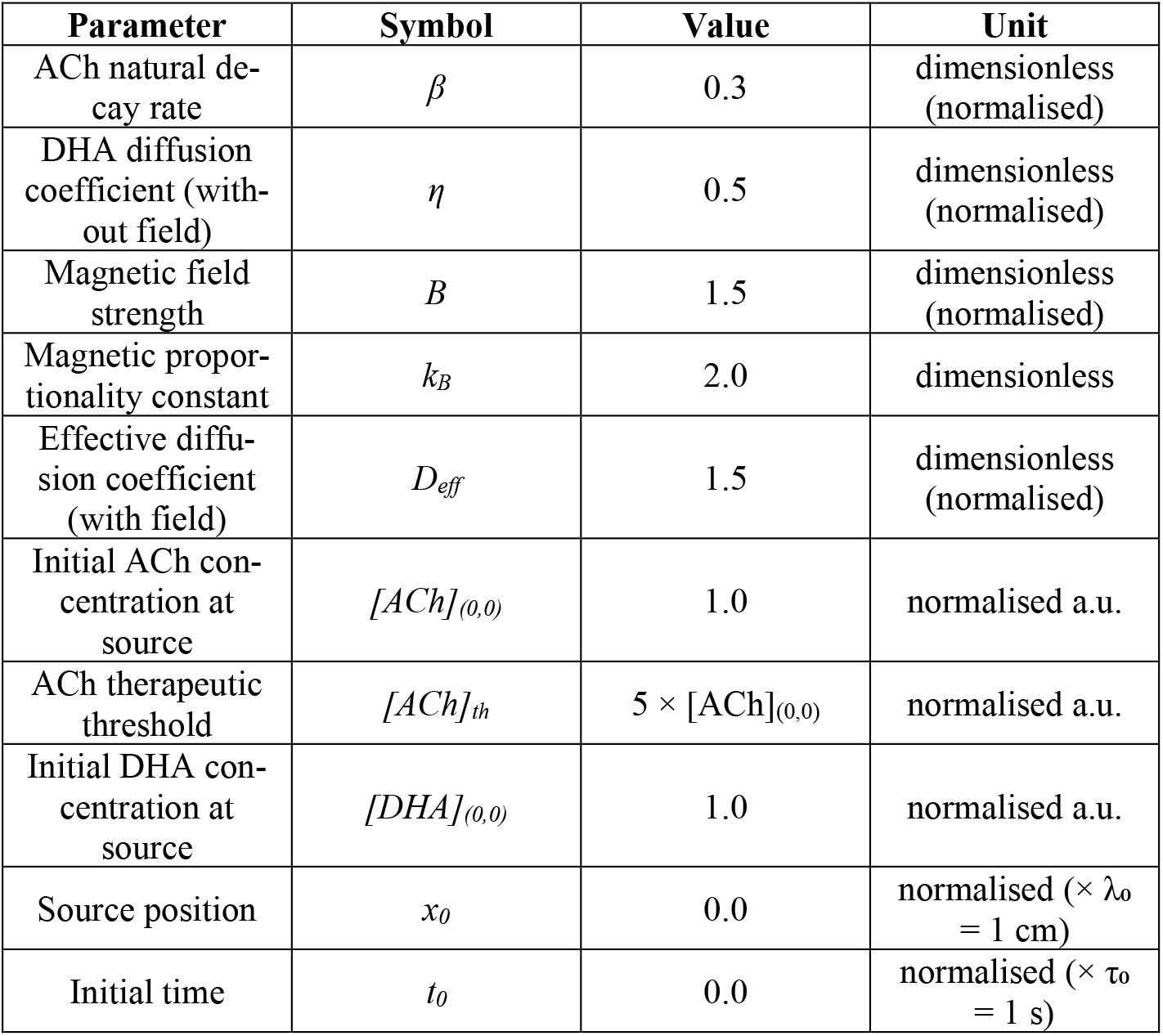
Model Parameters.

The coupling constant *α* from Equation (1) does not appear as an independent input. Under the effective-rate closure of Section 2.2, it is absorbed into the synthesis rate, which is modelled as equal to the diffusion coefficient, *η*. With the values in Table 1 (*k*_*B*_ *= 2*.*0, B = 1*.*5*), the effective diffusion coefficient evaluates to *D*_*eff*_ *= 1*.*5*. This is a threefold enhancement over the baseline value *η = 0*.*5*.

## 3 Simulation Methodology

The mathematical framework of Section 2 yields closed-form analytical expressions for all quantities of interest. As a result, the simulation requires no numerical integration of the governing equations, and no iterative ordinary-differential-equation solver. Instead, the DHA and ACh concentrations and the threshold-attainment times are obtained by directly evaluating the closed-form expressions, Equations (3)–(10), over the spatial and temporal domains of interest. All reported quantities are therefore analytic evaluations of the closed-form expressions, not numerical approximations. The computations and figures were implemented in Python. NumPy was used for array evaluation, and Matplotlib for figure generation.

### 3.1 Computational Procedure

The closed-form expressions are evaluated for two conditions throughout. The baseline case uses the diffusion coefficient, *η*. The magnetically enhanced case uses the effective coefficient, *D*_*eff*_. The threshold-time expressions [Equations (5) and (9)] give the time explicitly as a function of distance. The threshold-crossing times are therefore computed directly from these formulae, with no numerical root-finding. Three sets of quantities are computed, corresponding to the three quantitative outputs of Section 4:

- ACh concentration as a function of time at the representative observation point, 0.5 cm, evaluated for both conditions, together with the threshold-crossing times obtained from Equations (5) and (9). This produces the ACh concentration dynamics of Section 4.1 (Fig. 1).
- Threshold-attainment time as a function of distance across the modelled range 0– 3.0 cm from the source, for both conditions, giving the spatial threshold analysis of Section 4.2 (Fig. 2).
- DHA and ACh concentrations as functions of time over the co-evolution window 0–2.0 s at 0.5 cm, for both conditions, giving the DHA–ACh co-evolution analysis of Section 4.3 (Fig. 3).

**Fig. 1.**
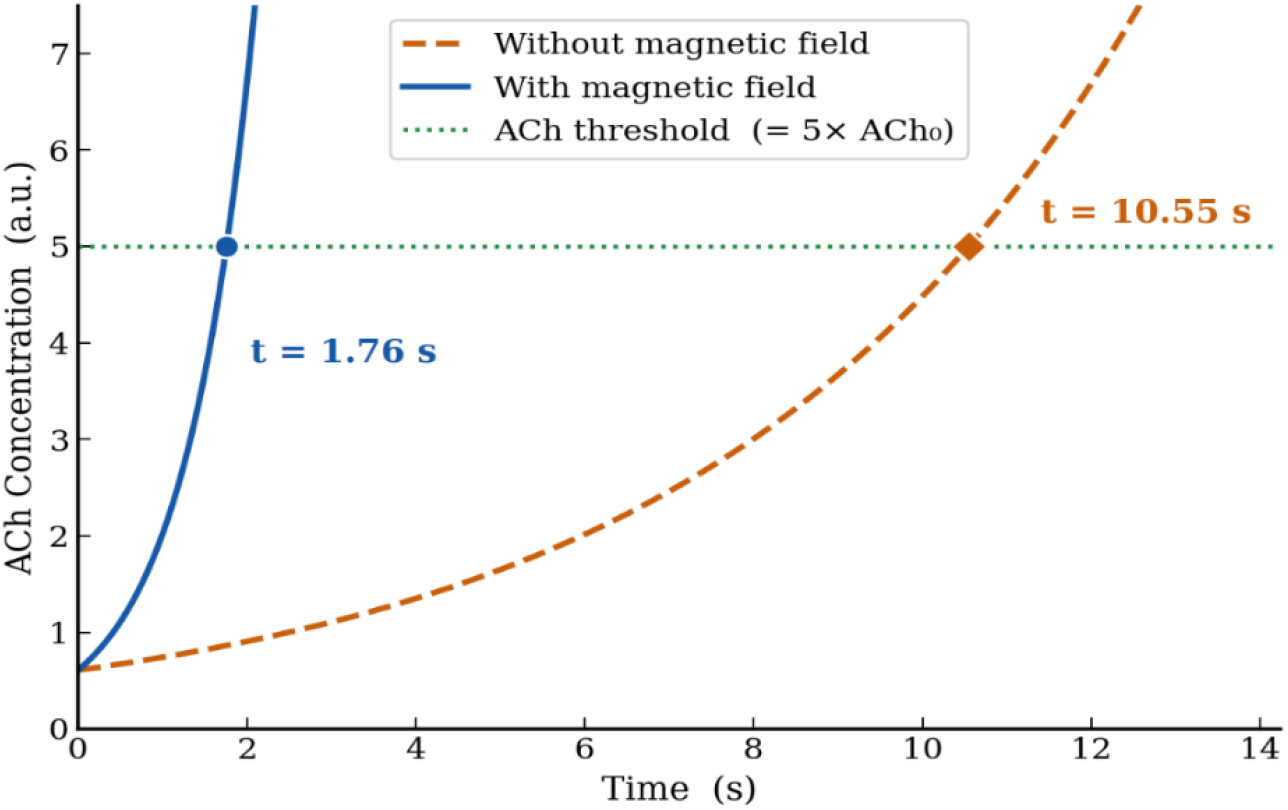
ACh concentration versus time at observation point, 0.5 cm from the DHA source.

**Fig. 2.**
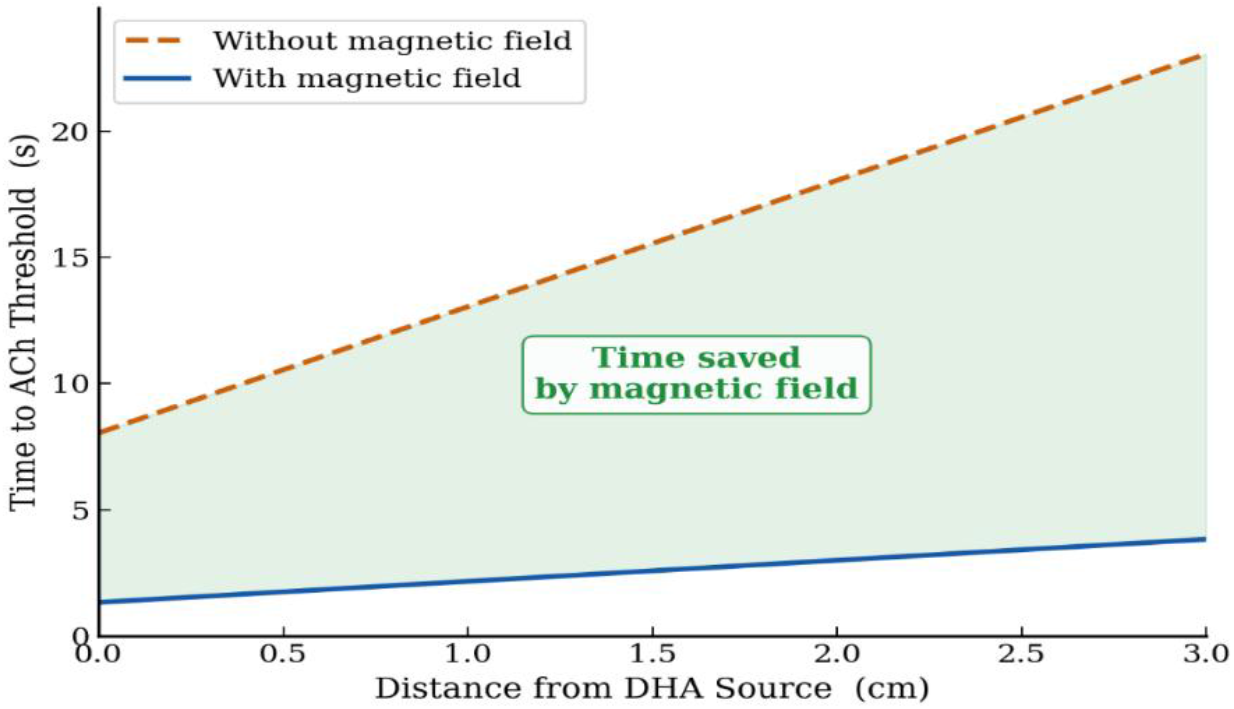
Time to reach the ACh threshold versus distance from the DHA source (0-3.0 cm).

**Fig. 3.**
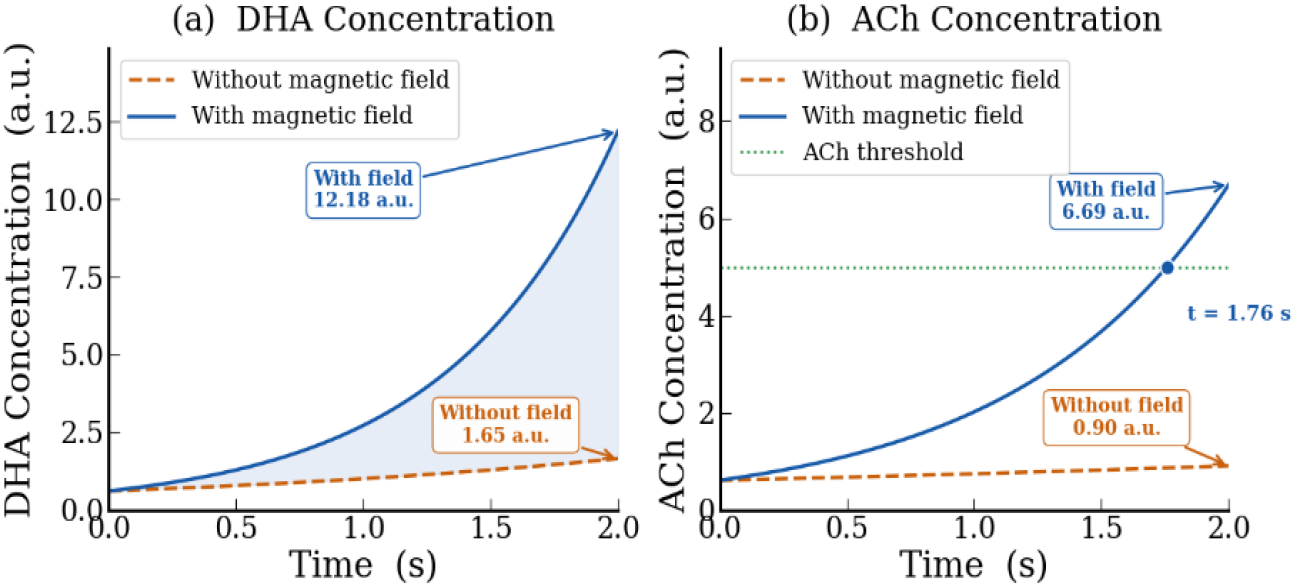
Co-evolution of DHA (left) and ACh (right) over time window 0-2.0 s at observation point, 0.5 cm.

The overall computational procedure is summarised in Algorithm 1. Each closed-form expression is sampled on a dense grid of several thousand points over the relevant domain for visualisation. The reported numerical values, for example, the threshold times of 10.55 s (baseline) and 1.76 s (with field) at observation point, 0.5 cm are computed directly from the closed-form expressions rather than read from the sampled grid. This ensures that all quoted figures are exact evaluations of the closed-form expressions.

#### Algorithm 1

Closed-Form Evaluation of DHA and ACh Dynamics

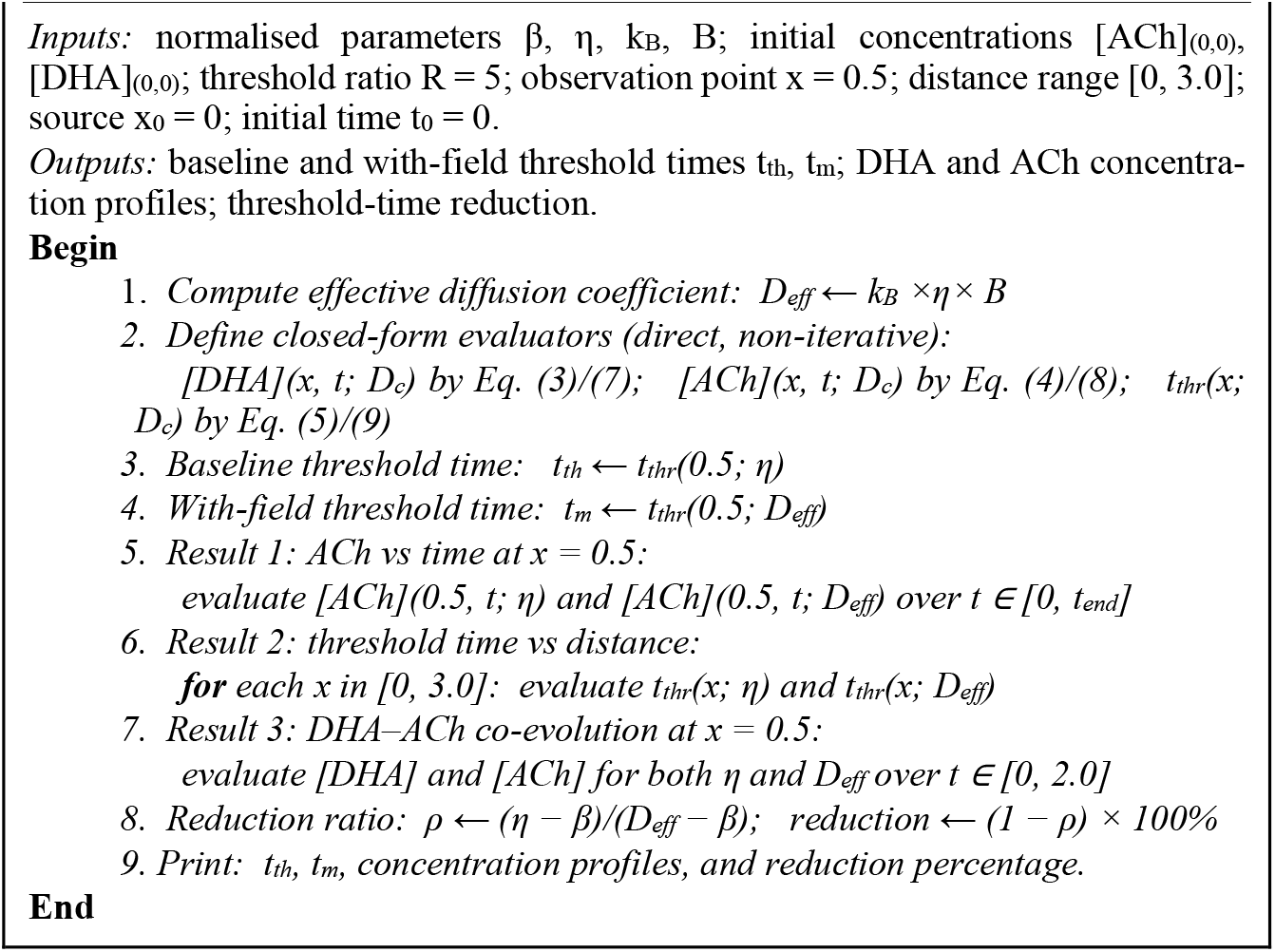

### 3.2 BCI-Based Closed-Loop Dosing Algorithm

The adaptive dosing framework is introduced as a future translational pathway (Section 4.4). It is implemented here as a rule-based, closed-loop simulation. A stimulus stream is generated over a 40 s window, sampled at 50 Hz. It follows an oddball paradigm, in which task-relevant target stimuli are interspersed among non-target stimuli. Two electrophysiological biomarkers are monitored within each target window. The first is the P300 event-related potential, which reflects cognitive target detection. The second is event-related desynchronisation/synchronisation (ERD/ERS), which reflects motor engagement.

The two biomarkers indicate the patient’s current cognitive–motor state. This inferred state determines the DHA dose, through three physiologically grounded states:

neither biomarker present → HIGH dose, indicating that maximal cholinergic support is required;
exactly one biomarker present (P300 only or ERD/ERS only) → MEDIUM dose;
both biomarkers present → ZERO dose, indicating that no additional support is required.

Dosing decisions are made only within the target-stimulus windows. The administered dose is thus adjusted in response to the patient’s real-time neurophysiological state. The dosing logic is summarised in Algorithm 2. This rule-based scheme provides the adaptive feedback mechanism that existing approaches lack (Section 1.2). Its demonstration across all four biomarker combinations, spanning the three dosing states is presented in Section 4.4.

#### Algorithm 2

BCI-Based Closed-Loop DHA Dose Regulation

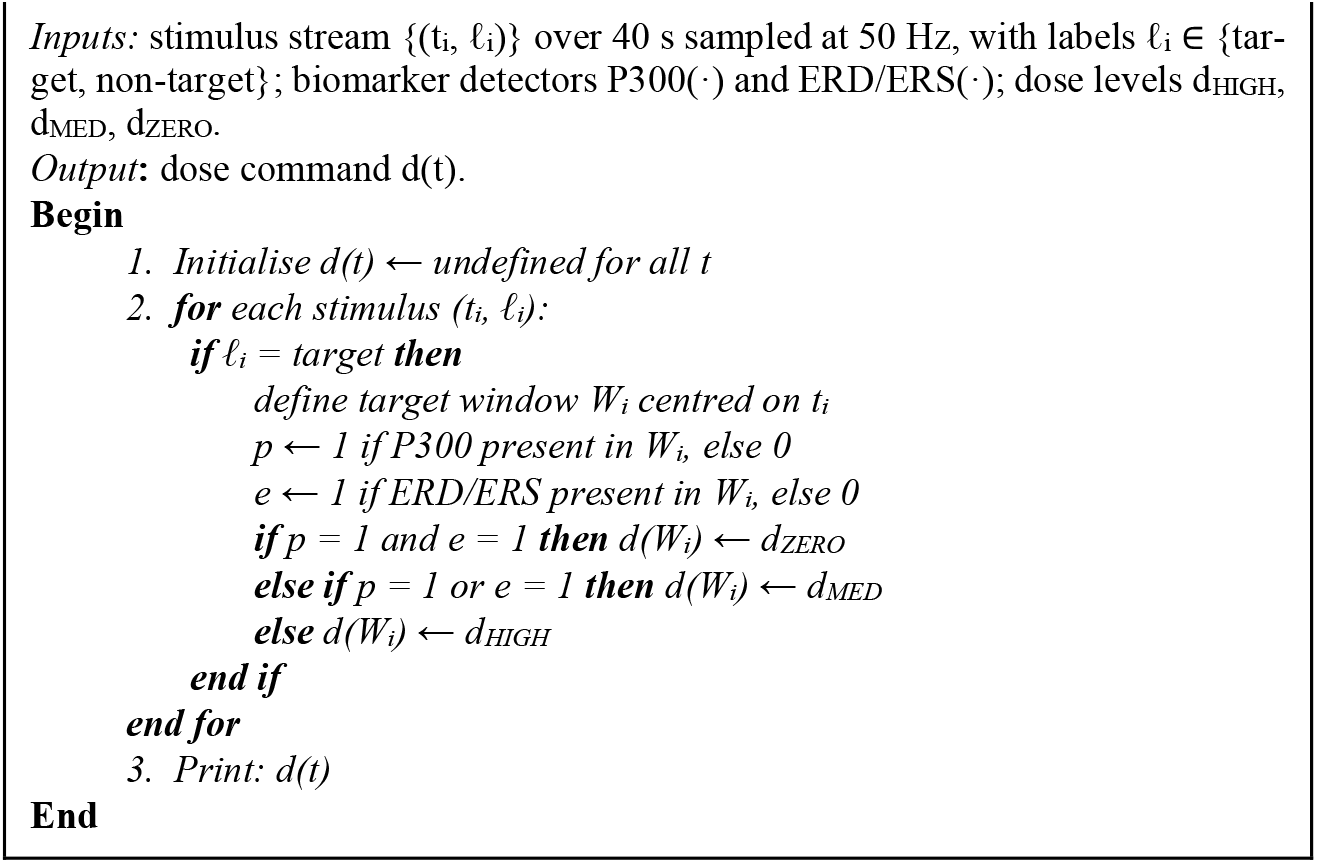

## 4 Results

The closed-form model of Section 2 was evaluated as described in Section 3. It produces three quantitative outputs. Each compares the baseline condition with the magnetically enhanced condition. A demonstration of the BCI-based closed-loop dosing framework follows. Time-course results are reported at the representative observation point, 0.5 cm. The spatial analysis spans 0-3.0 cm from the DHA source.

### 4.1 ACh Concentration Dynamics

Fig. 1 shows how ACh concentration evolves over time at the observation point, with and without magnetic field assistance. Both curves begin at the same initial concentration. The field therefore affects the rate at which ACh accumulates, not its starting level. Without the field, ACh reaches the therapeutic threshold in 10.55 s. With the field, it does so in only 1.76 s. This is an 83.3% reduction, corresponding to more than a five-fold acceleration. After crossing the threshold, the with-field curve continues to rise steeply. The benefit is therefore sustained, rather than confined to the initial phase. In the context of AD, reaching therapeutic ACh levels more rapidly implies a quicker cognitive benefit and a wider therapeutic window.

### 4.2 Spatial Threshold Analysis

Fig. 2 presents the threshold-attainment time as a function of distance from the source. In both conditions, the time increases linearly with distance. Across the full range, the with-field curve lies consistently below the baseline. The shaded band between them is the time saved by the field. The absolute saving widens with distance from 6.71 s at the source to 19.21 s at 3.0 cm. The proportional reduction, however, remains constant at 83.3% throughout rather than confined to the region near the source. The enhancement is therefore relevant to the range of brain targets affected in AD. Representative values are listed in Table 2.

**Table 2.** Threshold-attainment time versus distance from the source.

| Distance (cm) | Without field (s) | With field (s) | Time saved (s) | Reduction (%) |
| --- | --- | --- | --- | --- |
| 0.0 | 8.05 | 1.34 | 6.71 | 83.3 |
| 0.5 | 10.55 | 1.76 | 8.79 | 83.3 |
| 1.0 | 13.05 | 2.17 | 10.87 | 83.3 |
| 1.5 | 15.55 | 2.59 | 12.96 | 83.3 |
| 2.0 | 18.05 | 3.01 | 15.04 | 83.3 |
| 2.5 | 20.55 | 3.42 | 17.12 | 83.3 |
| 3.0 | 23.05 | 3.84 | 19.21 | 83.3 |

### 4.3 DHA-ACh Co-evolution

Fig. 3 shows the joint evolution of DHA and ACh over the 0-2.0 s window at the observation point. In the left panel, DHA accumulates far faster under field assistance. It reaches 12.18 a.u at 2 s, compared with 1.65 a.u without the field. This is a 7.39-fold increase. In the right panel, the greater DHA availability drives ACh upward more rapidly. ACh crosses the therapeutic threshold at 1.76 s and reaches 133.7% of it by 2 s. Without the field, it reaches only 18.1%. These results make the underlying causal chain explicit. Greater DHA delivery raises the DHA concentration at the target. This accelerates ACh synthesis, which in turn brings the threshold crossing forward. The field does not merely speed up DHA delivery. It keeps DHA elevated throughout the window, so ACh is driven both sooner and to a higher level.

### 4.4 BCI-Based Closed-Loop Dosing Framework

Fig. 4 demonstrates the closed-loop dosing framework (Algorithm 2) on a 40 s stimulus stream. The stream contains four target events. These were selected to exercise all four combinations of the two biomarkers, and hence all three dosing states. For the first target, neither the P300 nor the ERD/ERS biomarker is detected, and the framework issues a HIGH dose. For the second target, only the P300 is present. For the third, only the ERD/ERS is present. In both cases a MEDIUM dose is issued. For the fourth target, both biomarkers are present, and a ZERO dose is issued. Dose commands are produced only within the target windows, and adapt to the inferred cognitive–motor state. No command is issued between targets.

**Fig. 4.**
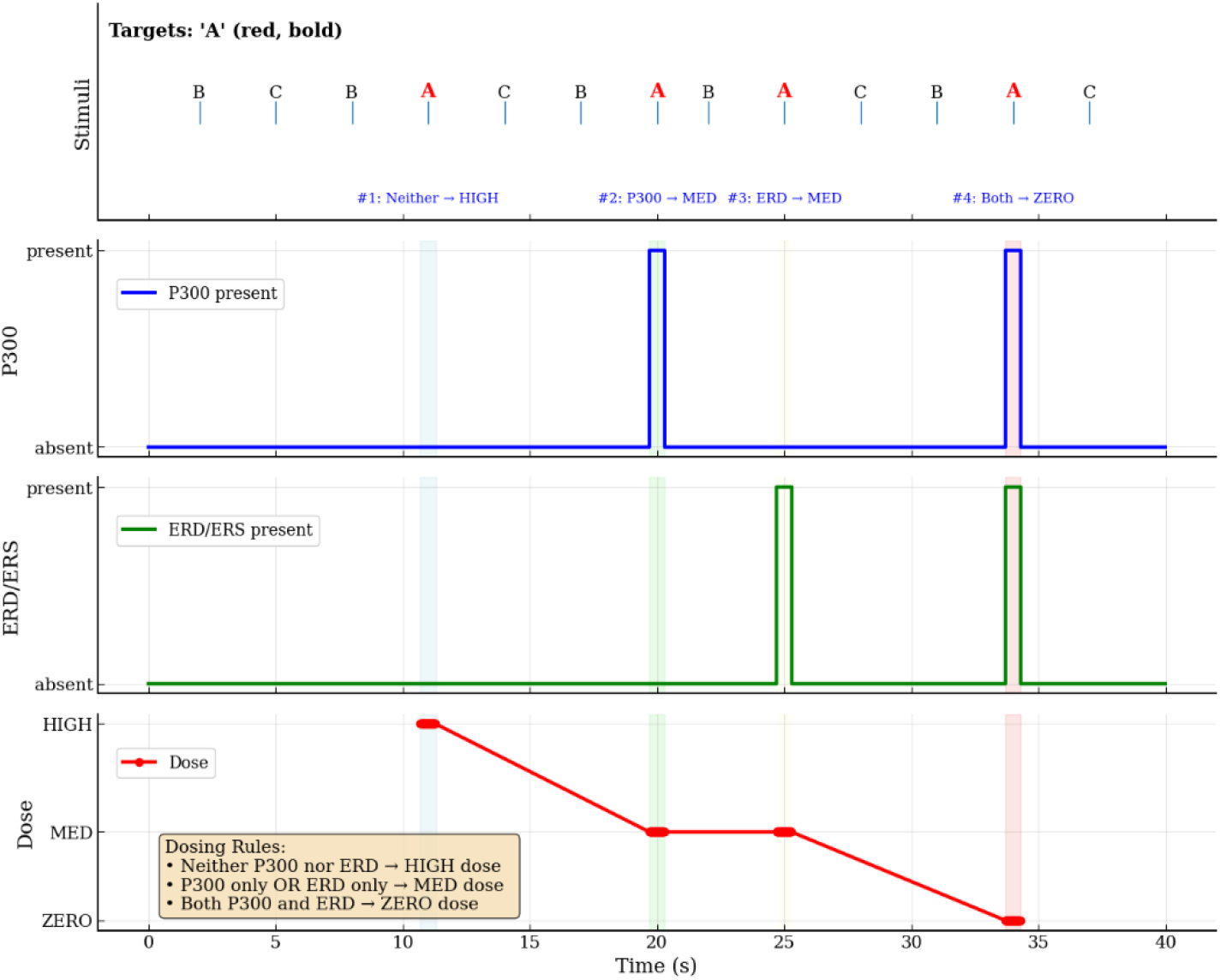
Demonstration of the BCI-based closed-loop dosing framework over a 40 s stimulus stream. From top to bottom: the stimulus sequence (targets marked A), P300 presence, ERD/ERS presence, and the resulting dose command.

This demonstrates closed-loop operation. The framework maps the momentary biomarker state to a graded dose. It thereby provides the adaptive feedback that open-loop supplementation lacks. The present result establishes the control logic as a proof of concept. Biomarker acquisition and dose-response calibration for clinical use are proposed as future work (Section 6).

## 5 Discussion

This study shows that a static magnetic field can speed up the restoration of ACh. The field acts on the natural diamagnetic anisotropy of DHA. The model was evaluated at an observation point, 0.5 cm from the DHA source, across a modelled range of 0–3.0 cm, and over a 2.0 s observation window. Three results stand out.

First, the field shortens the time needed for ACh to reach its therapeutic threshold. At the observation point, this time falls from 10.55 s to 1.76 s, a reduction of 83.3%. Second, this benefit holds right across the modelled range. The absolute time saved grows with distance, from 6.71 s at the source to 19.21 s at the far end of the range. The percentage reduction, however, stays fixed at 83.3%. However, it is worth mentioning that the uniformity is a consequence of how the model is built, not a separate empirical finding. What the model does show is that the benefit does not fade with depth. This matters in practice, because the brain regions affected in AD lie at different depths. Third, the co-evolution analysis explains the cause. The field raises the effective diffusion coefficient of DHA. As a result, by the end of the 2.0 s window, the DHA level at the target is 7.39 times its baseline value. This drives ACh to 133.7% of the threshold, compared with only 18.1% without the field.

The biological assumptions of the model are supported by the previous work. Dietary DHA raises cerebral ACh (Minami *et al*. 1997). DHA also affects neurotransmitter release and synaptic function (Vancassel *et al*. 2007). A lack of DHA lowers muscarinic receptor binding (Aïd *et al*. 2003). These studies establish a link between DHA and cholinergic signalling. The present model turns that qualitative link into an explicit description of how the two concentrations change over time.

The magnetic mechanism builds on earlier biophysical work. Rosen (2003) examined how moderate static magnetic fields affect biological systems. He found that such fields can alter the behaviour of membrane ion channels. To explain this, he pointed to a property called diamagnetic anisotropy. Molecules with this property reorient when a magnetic field is applied. In Rosen’s account, the molecules were the phospholipids of the cell membrane. DHA is one of the fatty acids carried by these membrane phospholipids, and it shares the same property: it is diamagnetically anisotropic.The present work uses this reorientation principle in a different way. Here it is applied to the movement of DHA through tissue. When DHA aligns with the field, its transport becomes more directional. This raises its effective diffusion coefficient. This step is proposed in the present work. It is not a finding of the cited study. The approach also differs from recent magnetic delivery methods that inject magnetic nanoparticles (Senanayake *et al*. 2025; Alipour *et al*. 2025). No added agents are needed here, because DHA already has the required property. To the authors’ knowledge, this is the first closed-form model to link magnetically enhanced DHA transport with the kinetics of ACh restoration.

These results suggest a different way of delivering DHA. Ordinary dietary supplementation is limited by poor passage across the blood–brain barrier and by slow diffusion (Pan *et al*. 2015; Iwao *et al*. 2023). Magnetic assistance offers an active and non-invasive alternative. The proposed wearable coils give a practical way to generate the field. The BCI-based closed-loop scheme adds a further step: it adjusts the dose to the patient’s current cognitive–motor state. Both remain proposals rather than tested components. Together, however, they point toward a form of DHA therapy that is actively controlled and responsive to the patient’s condition. The same approach may, in principle, suit other disorders in which cholinergic signalling is impaired. These conclusions hold within the assumptions of the model. The closed-form results are exact for the linearised system, and all quantities are normalised, so they describe relative acceleration rather than clinical values. The limitations that follow are considered in Section 6.

## 6 Limitations and future experimental directions

The present study is an *in silico* investigation, and its conclusions hold within the assumptions of the model.

### 6.1 Limitations

The first is the linearising closure. The synthesis rate *α×[DHA]* is modelled as an effective rate constant equal to *η* (Section 2.2). This makes the analysis tractable, but it means that concentration expressions and threshold-time expressions are exact for the linearised model rather than for the full nonlinear system. Moreover, the magnitude of the predicted reduction in threshold time depends on the assumed magnetic parameters. However, the spatial uniformity of the reduction follows from the structure of the model, not from any particular parameter choice.

Secondly, the study is entirely in silico, and expressed in normalised units. The results therefore describe the relative speed-up of ACh restoration and how it scales with distance. They are not predictions of dimensional clinical values, and they have not yet been tested experimentally. Transport is also modelled along a single spatial coordinate so the spatial predictions are simplified.

Lastly, the physical mechanism (field-enhanced DHA transport) and the delivery components (wearable coils and BCI dosing scheme) are proposed rather than validated. Previous work (Rosen, 2003) attributes the effects of static magnetic fields to the reorientation of diamagnetically anisotropic membrane phospholipids. The present work extends that principle to the transport of DHA itself, which requires experimental confirmation. The wearable coil configuration and the BCI-based dosing scheme are likewise design proposals.

### 6.2 Future Work

Experimental validation is the most immediate priority. Microdialysis-based measurement of extracellular ACh, with and without an applied field would test the predicted time course directly. It would also supply the parameter values needed to express the model in physical units, and would confirm whether a static field enhances DHA transport as assumed.

The model can also be extended. The nonlinear system can be studied and the single spatial coordinate can be replaced by a realistic geometry.

Finally, the translational components require development. A wearable coil would need design, field characterisation and safety assessment. The BCI scheme would need real-time biomarker acquisition from recorded EEG, followed by dose–response calibration. These steps are necessary before any clinical evaluation could be considered. The same approach may in principle suit other disorders of cholinergic signalling, which remains to be examined.

## 7 Conclusion

This paper has presented a closed-form model of magnetic field-enhanced DHA delivery for ACh restoration in AD. The model couples DHA diffusion with ACh synthesis. It yields explicit expressions for both concentrations, and for the time to reach a therapeutic threshold, with and without an applied static magnetic field.

Evaluation of the model shows that field assistance shortens the threshold time by 83.3% at the observation point. The proportional reduction holds across the modelled range, and the co-evolution analysis identifies raised DHA availability as its cause. Together these results give a quantitative basis for treating DHA delivery as an actively controlled process rather than a passive one. A BCI-based closed-loop dosing scheme is also demonstrated, as a route toward dosing that responds to the patient’s current state.

The findings are those of a model, and hold within its assumptions. Experimental measurement of ACh under an applied field is the necessary next step, together with the extensions set out in Section 6.

## References

1. Aïd, S., Vancassel, S., Poumès-Ballihaut, C., Chalon, S., Guesnet, P. and Lavialle, M. (2003). Effect of a diet-induced n-3 PUFA depletion on cholinergic parameters in the rat hippocampus. Journal of Lipid Research,.44, pp. 1545–1551.

2. Ali, A.H., Hachem, M. and Ahmmed, M.K. (2024) Docosahexaenoic acid-loaded nanoparticles: A state-of-the-art of preparation methods, characterization, functionality, and therapeutic applications, Heliyon, 10, e30946. doi:10.1016/j.heliyon.2024.e30946.

3. Alipour Mozhgan, Abdolmaleki Maryam, Shabanpour Yaser, Zali Alireza, Ashrafi Farzad, Nohesara Shabnam, Hajipour-Verdom Behnam, Advances in magnetic field approaches for non-invasive targeting neuromodulation, Frontiers in Human Neuroscience, 19 (2025), https://www.frontiersin.org/journals/human-neuroscience/articles/10.3389/fnhum.2025.1489940, DOI=10.3389/fnhum.2025.1489940, ISSN=1662-5161.

4. Birks, J. S., & Harvey, R. J. (2018). Donepezil for dementia due to Alzheimer’s disease. The Cochrane database of systematic reviews, 6(6), CD001190. 10.1002/14651858.CD001190.pub3.

5. Cummings, J., Lee, G., Zhong, K., Fonseca, J. and Taghva, K. (2021) Alzheimer’s disease drug development pipeline: 2021, Alzheimer’s & Dementia: Translational Research & Clinical Interventions, 7(1), e12179.

6. Duro, M. V. V., Van Valkenburgh, J., Ingles, D. E., Tran, J., Cai, Z., Ebright, B., Wang, S., Kerman, B. E., Galvan, J., Hwang, S. H., Sta Maria, N. S., Zanderigo, F., Croteau, E., Cunnane, S. C., Rapoport, S. I., Louie, S. G., Jacobs, R. E., Yassine, H. N., & Chen, K. (2023). Synthesis and Preclinical Evaluation of 22-[18F] Fluorodocosahexaenoic Acid as a Positron Emission Tomography Probe for Monitoring Brain Docosahexaenoic Acid Uptake Kinetics. ACS chemical neuroscience, 14(24), 4409–4418. 10.1021/acschemneuro.3c00681.

7. Francis, P.T., Palmer, A. M., Snape, M., Wilcock, G.K. (1999). The cholinergic hypothesis of Alzheimer’s Disease: a review of progress. Journal of Neurology: Neurosurgery and Psychiatry, 66, pp. 137–147.

8. Hachem M, Belkouch M, Lo Van A, Picq M, Bernoud-Hubac N, Lagarde M. Brain targeting with docosahexaenoic acid as a prospective therapy for neurodegenerative diseases and its passage across blood brain barrier. Biochimie. 2020 Mar;170:203–211. doi: 10.1016/j.biochi.2020.01.013. Epub 2020 Jan 31. PMID: 32014503.

9. Iwao, T., Takata, F., Matsumoto, J., Aridome, H., Yasunaga, M., Yokoya, M., Kataoka, Y., & Dohgu, S. (2023). Aging decreases docosahexaenoic acid transport across the blood-brain barrier in C57BL/6J mice. PloS one, 18(2), e0281946. 10.1371/journal.pone.0281946.

10. Kasa, P., Rakonczay, Z., Gulya, K. (1997). The cholinergic system in Alzheimer’s Disease. Progress in Neurobiology, 52, pp. 511–535.

11. Kim, H. Y., Huang, B. X., & Spector, A. A. (2022). Molecular and Signaling Mechanisms for Docosahexaenoic Acid-Derived Neurodevelopment and Neuroprotection. International journal of molecular sciences, 23(9), 4635. 10.3390/ijms23094635.

12. Minami, M., Kimura, S., Endo, T., Hamaue, N., Hirafuji, M., Togashi, H., Matsumoto, M., Yoshioka, M., Saito, H., Watanabe, S., Kobayashi, T., Okuyama, H. (1997). Dietary Docosahexaenoic Acid Increases Cerebral Acetylcholine Levels and Improves Passive Avoidance Performance in Stroke-Prone Spontaneously Hypertensive Rats. Pharmacology Biochemistry and Behaviour, 58(4), pp. 1123 1129.

13. Pan, Y., Khalil, H., & Nicolazzo, J. A. (2015). The Impact of Docosahexaenoic Acid on Alzheimer’s Disease: Is There a Role of the Blood-Brain Barrier?. Current clinical pharmacology, 10(3), 222–241. 10.2174/157488471003150820151532.

14. Rapoport, S.I., Ramadan, E. and Basselin, M. (2011). Docosahexaenoic acid (DHA) incorporation into the brain from plasma, as an in vivo biomarker of brain DHA metabolism and neurotransmission. Prostaglandins & Other Lipid Mediators, 96 (14), pp. 109–113.

15. Rosen, A.D. (2003). Mechanism of action of moderate-intensity static magnetic fields on biological systems. Cell Biochemistry and Biophysics, 39(2), pp. 163–173.

16. Senanayake D, Yapa P, Dabare S, Munaweera I. Precision targeting of the CNS: recent progress in brain-directed nanodrug delivery. RSC Adv. 2025 Jul 21;15(32):25910–25928. doi: 10.1039/d5ra03578c. PMID: 40697447; PMCID: PMC12278267.

17. Singh, M. (2005). Essential Fatty Acids, DHA and Human Brain. Indian Journal of Pediatrics, 72, pp. 239–242.

18. Tanaka, K., Farooqui, A.A., Siddiqi, N.J., Alhomida, A.S. and Ong, W. (2012). Effects of Docosahexaenoic Acid on Neurotransmission. Biomolecules and Therapeutics, 20(2), pp. 152–157.

19. Vancassel, S., Aid, S., Denis, I., Guesnet, P. and Lavialle, M. (2007). DHA involvement in neurotransmission process. Journal of Oilseeds and fats Crops and Lipids, 14 (3-4), pp. 164–170.

20. Weiser, M.J., Butt, C.M., Mohajeri, M.H. (2016). Docosahexaenoic Acid and Cognition throughout the Lifespan. Nutrients, 8, 99. doi:10.3390/nu8020099.

21. Yiannopoulou, K. G., & Papageorgiou, S. G. (2020). Current and Future Treatments in Alzheimer Disease: An Update. Journal of central nervous system disease, 12, 1179573520907397. 10.1177/1179573520907397.

